# Insecticide temephos alters thermal dependence of dengue vector

**DOI:** 10.64898/2026.04.01.715840

**Authors:** Patrick M. Heffernan, Courtney C. Murdock, Jason R. Rohr

## Abstract

1. Although ecological research has long focused on the effects of temperature on population growth, arthropod pests are exposed to a wide variety of environmental factors that affect their performance, such as chemical pesticides targeted against them. Moreover, these environmental factors likely do not act in isolation. Identifying the extent to which abiotic factors interact to affect pest population dynamics can strengthen current and future pest management programs.
2. Here, we investigated the extent to which temephos, a common pesticide applied to aquatic environments for mosquito control, influences the thermal performance of juvenile survival and development rate, as well as the intrinsic population growth rate, of the invasive mosquito pest, *Aedes aegypti.* We implemented a response surface experimental design to measure these traits across seven temperatures and five temephos concentrations and fit temperature- and insecticide-dependent performance curves to assess impacts on the overall performance and the thermal optimum, minimum, and maximum.
3. Temephos exposure profoundly altered the thermal performance of juvenile survival by reducing survival across all temperatures, shrinking the thermal breadth, and shifting the thermal optimum to warmer temperatures. Through this, temephos also altered the thermal performance of population growth primarily by reducing its thermal breadth.
4. *Synthesis and applications:* Our findings demonstrate that interactions between temperature and insecticide exposure can fundamentally reshape pest population dynamics, rather than acting as independent stressors. By quantifying this interaction, we showed that temphos is most effective below the pest’s thermal optimum, suggesting that larvicides may yield the greatest population suppression in cooler regions or during cooler periods of the year. Incorporating such temperature-dependent efficacy into pest management strategies could improve the timing and spatial targeting of control efforts. More broadly, these results highlight the need to integrate anthropogenic stressors with climatic drivers when predicting pest risk and optimizing management under ongoing environmental change.

## Introduction

Arthropod pests pose major and growing challenges to human health and agricultural production worldwide. Vector-borne diseases cause over 700,000 human deaths annually and impose economic costs estimated in the hundreds of billions of US dollars (Bradshaw et al., 2016; Roiz et al., 2024). Agricultural pests similarly cause substantial crop destruction and livestock production losses annually (Culliney, 2014; Narladkar, 2018; Taylor et al., 2012). Optimizing pest control under accelerating environmental change requires understanding how multiple environmental drivers jointly influence population dynamics.

Because arthropods are ectothermic, ecological research has long emphasized how environmental temperature shapes their activity, abundance, and geographic distribution (Abram et al., 2017; Kingsolver & Buckley, 2020; Nascimento et al., 2022; Sinclair et al., 2003). Consistent with the Metabolic Theory of Ecology, temperature effects on biological traits are often nonlinear, with performance following unimodal, hump-shaped relationships known as thermal performance curves (TPCs) (Arnoldi et al., 2025; Mordecai et al., 2019; Pawar et al., 2024). Trait TPCs are characterized by lower and upper limits (*T_min_* and *T_max_*) and an optimal temperature (*T_opt_*) at which performance is maximized. Incorporating trait TPCs into temperature-dependent population-level parameters, such as intrinsic growth rate (*r_m_*) and *R_θ_*, has improved predictions of pest distributions and dynamics across geographic and temporal scales, particularly for mosquitoes (Mordecai et al., 2019). These temperature-dependent frameworks can be used to forecast pest risk and guide intervention strategies (Mordecai et al., 2019, 2020; Ryan et al., 2019a).

Insecticides represent one of the most widely deployed tools for pest suppression. However, their effects are rarely experienced as static or uniformly lethal in natural systems. Chemical degradation, heterogeneous application, and environmental variation in bioavailability frequently result in sublethal exposures (Andreazza et al., 2021; Guedes et al., 2017; Müller, 2018). In mosquitoes, sublethal concentrations have been shown to influence all life history traits underlying *r_m_*, such as mortality, fecundity and development rate (Alvarez Costa et al., 2018; Muturi et al., 2011; Pelizza et al., 2010; Reyes-Villanueva et al., 1990, 1992; Shaalan et al., 2005). Despite these documented trait-level effects, population models typically treat insecticide impacts as temperature-invariant or fail to incorporate their full suite of trait-mediated consequences (Barbosa et al., 2018; Elliott et al., 2018; Ngonghala et al., 2021). This limits our ability to predict when and where chemical control will be most effective.

Critically, insecticide efficacy is temperature dependent (Moyes et al., 2017). Mosquito larvae, for example, can mitigate larvicide exposure through metabolic resistance, sequestering and detoxifying toxins in aquatic environments (Li et al., 2007; Liu, 2015; Moyes et al., 2017). Both metabolic rates and insecticide toxicity vary with temperature, suggesting that chemical control may reshape TPCs rather than simply reduce overall performance. Nguyen et al. (2021) used model simulations in the schistosomiasis system to demonstrate that control measures targeting intermediate hosts of pathogens can shift the *T_opt_* for disease transmission, altering both predicted disease distributions and temperature at which control is most effective. Whether similar temperature-dependent shifts occur in insecticide-driven pest systems remains unknown.

In this study, we test whether a commonly used larvicide, temephos, alters the thermal performance (optimum, minimum, and maximum) of juvenile survival, development rate, and intrinsic population growth rate (*r_m_*) in the invasive mosquito *Aedes aegypti*. More than 6 billion people are at risk of viral diseases vectored by *Ae. aegypti*, including dengue, Zika, chikungunya, or yellow fever, and both geographic risk and insecticide use are expected to increase over the coming decades due to environmental changes (Ryan et al., 2019b; Shattuck et al., 2023). We hypothesized that sublethal temephos exposure would interact with temperature to (i) modify the thermal optima and limits of juvenile survival and development, and (ii) shift or compress the thermal performance of *rₘ*.

To test these predictions, we conducted a response surface experiment spanning seven temperatures (17°C-41°C) and five temephos concentrations (0-0.0127 ppm) on juvenile *Ae. aegypti* mosquitoes (Table S1). Across these gradients, we quantified survival to adult emergence, development rate, and wing length at emergence, and used these trait responses to calculate *r_m_.* We then mapped *r_m_* globally using seasonal temperatures to demonstrate the extent to which pesticide exposure can shape thermal predictions. By integrating chemical exposure into a temperature-dependent population framework, this study advances our ability to predict how environmental heterogeneity shapes pest control outcomes and identifies conditions under which larvicides are likely to be most effective.

## Methods

### Maintenance of field collected *Aedes aegypti* Thai strain and resistance profile

A field-collected strain (Nakhon Ratchasima Province, Thailand, by Alongkot Ponlawat, and shared by Courtney Murdock, Cornell University) was maintained in standard rearing conditions (27°C, 80% RH ± 10% RH, 12:12 light-dark cycle) for five generations. To avoid effects of crowding, 50 larvae were sorted into 450 mL of water and fed one fish food pellet (Cichlid Gold, Hikari, Hayward, USA) every three days or *ad libitum.* Adult mosquitoes were provided a human blood meal via a water-jacketed membrane feeder twice a week and were also supplied with a 10% sucrose solution via soaked cotton balls. Prior to the study, resistance to temephos was characterized using a WHO dose-response assay kit (WHO VBC, 1981). The LC50 of the F5 *Ae. aegypti* Thai strain was determined to be 0.014 ppm (SE = 0.0006).

### Experimental design

We tested a response surface of seven constant temperatures (17°C to 41°C in 4°C increments) and five temephos concentrations, totaling 35 unique treatment combinations. To increase feasibility, we utilized the response surface to remove eight treatment combinations (Inouye, 2001), reducing the number of unique combinations from 35 to 27 (see Table S1). Temperature was maintained using 27 independent incubators set to target temperatures (incubators described in Raffel et al. [2013]). The actual temperature was measured at the base of each rearing cup daily using an infrared thermometer (Etekcity Lasergrip 1080, Vesync Group, Shenzhen, China), and a temperature logger (Onset HOBO pendant data logger, Onset Computer Corporation, Pocasset, USA) was kept on each incubator’s heating apparatus to measure temperature consistency throughout the course of the experiment. To ensure only stable incubators were used in the fitting of TPCs, any incubator with an actual temperature standard deviation ≥2°C was removed. Furthermore, any incubators whose mean actual temperatures differed from their target setpoint were treated as the nearest target temperature treatment in the analyses. We chose to bin the temperatures rather than use the exact mean to allow for adequate replication at each discrete temperature point to characterize the mean response. In the first temporal block, we tested five temephos concentrations (technical grade temephos provided by BASF; dissolved in 200-proof ethanol) with the LC50 of the strain as the maximum concentration (0 ppm, 0.008 ppm, 0.01 ppm, 0.012 ppm, and 0.014 ppm). After this temporal block was completed, the concentrations were adjusted so the LC40 (0.0127 ppm) was the maximum concentration for three more temporal blocks (0 ppm, 0.007 ppm, 0.009 ppm, 0.0111 ppm, and 0.0127 ppm). In the analyses, results from the first temporal block were treated as the nearest concentration tested in the second through fourth temporal blocks. To ensure this merging did not bias the trait fits, we conducted the TPC analyses (see *Thermal performance curve fitting*) on both all temporal blocks and just the second through fourth temporal blocks. We found that the parameter estimates did not differ between these two analyses (assessed through overlapping 95% credible intervals).

Within each incubator, 25 1st instar larvae were sorted into a styrofoam treatment cup with 200 mL of water and ample food available within each incubator. 48 hours after hatching, temephos was added to each cup and gently mixed. In total, 2,700 juvenile mosquitoes were tested (2,025 across the second through fourth temporal blocks and 675 from the first temporal block). Until all immature mosquitoes had emerged as adults or died, cups were monitored daily for pupation. Upon pupation, pupae were removed from the rearing cup and placed in 50 mL conical centrifuge tubes with 10 mL of water within the incubator to track time from pupation to emergence. Survival probability was recorded as the proportion of mosquitoes surviving to adult emergence from each cup, and development rate was recorded as the inverse of time from egg to adult emergence.

After emergence, adult mosquitoes were removed from the incubators and euthanized in a freezer. Once euthanized, the mosquitoes were identified as male or female, and one wing per mosquito was removed and imaged under a dissecting scope. The wing length (mm) of each individual was measured using Image-J (Schneider et al., 2012).

### Thermal performance curve fitting

To assess how *T_opt_, T_min_,* and *T_max_* of juvenile survival probability and development rate changed with temephos exposure, we fit Bayesian TPCs using *R2Jags* (Su et al., 2015) in R version 4.5.1 (R Core Team, 2025). For each trait, we fit quadratic and Brière curves in which the *T_min_* and *T_max_* parameters were monotonically dependent on temephos concentration (adapted from Dennington et al [2024]). For full details of the model formulas and priors, see the *Supporting Information*. Each curve was fitted using 3 MCMC chains, 50,000 burn-in iterations, 450,000 saved iterations, and a thinning rate of 15. Quality of each fit was assessed through the Gelman-Rubin statistic, effective sample size, and trace plots.

### Wing length of adults

To assess the interaction of temperature and temephos concentration on mosquito wing length, we fit a generalized linear mixed effects model (GLMM) using the *glmmTMB* (Brooks et al., 2017) package in R version 4.5.1 (R Core Team, 2025). In this GLMM, we included main effects for temperature and temephos concentration as well as their interaction. We also included a random effect for larval rearing cup to account for cup-specific differences in wing length. We conducted these analyses separately on all emerged mosquitoes and just the female emerged mosquitoes.

### Intrinsic growth rate

To estimate *r_m_* across temperature and temephos concentrations, we used a derivation of the Euler-Lotka framework applied to arthropods from Huxley et al (pre-print, 2025) and Cator et al (2020) (equation 1).

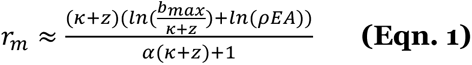

In equation 1, *z* is adult mortality rate, *b_max_* is maximum reproductive rate (eggs per adult female per day), 𝜅 is is fecundity loss (per individual per day), 𝜌𝐸𝐴 is survival probability from egg to adulthood, and 𝛼 is egg-to-adulthood development time (days). As we did not measure how temperature and temephos interact to affect adult life history traits, we used constant values of 0.041 days^−1^ for mortality rate (*z*) and 3.309 eggs per female per day (*b_max_*) based on the mean values of these traits from the *AedesTraits* database (Huxley et al., 2025).

### Sensitivity analyses

We performed sensitivity analyses to determine the relative contributions of juvenile survival probability and development rate on *r_m_* across temperatures and temephos concentrations. In line with the chain rule method applied by Shocket et al (2020) and Huxley et al (pre-print, 2025), at each temephos concentration, we calculated the derivatives of each trait across temperatures and then multiplied them by the partial derivatives of *r_m_* with respect to each trait (while keeping the non-derived trait at their mean) across temperatures. We then plotted the sensitivity of *r_m_* and the relative sensitivities of juvenile survival probability and development rate across temperatures and temephos concentrations (Fig. S1). Results of the sensitivity analyses are reported in the *Supporting Information*.

### Mapping thermal suitability in the context of varying insecticide exposure

To demonstrate how we may expect low concentrations of temephos to shift the *Aedes aegypti* population landscape, we used ERA5 monthly land temperature (Muñoz Sabater, 2019) from 2024 to map population growth in three temephos exposure scenarios (0 ppm, 0.009 ppm, and 0.0127 ppm). We first calculated the mean monthly temperature across the globe for four quarters of 2024 (January-March, April-June, July-September, and October-December) using Google Earth Engine (Gorelick et al., 2017). We then used the *terra* package (Hijmans et al., 2026) in R to map each *r_m_* curve’s median values to the mean temperature values. As there is no global dataset detailing where and to what degree mosquito populations across the globe are being exposed to temephos, these maps are a heuristic tool to explore potential implications for variation in insecticide exposure on thermal suitability and are not intended to be used for predictions. Rather, this mapping exercise is an exploration of the degree to which temephos has the potential to shift mosquito population dynamics and highlight regions in which temephos may be most and least effective at reducing population growth.

## Results

### Effects of temperature and temephos concentration on juvenile survival probability

The thermal response of juvenile survival probability showed strong dependencies on temephos concentration (Fig. 1A). Non-overlapping 95% highest posterior density (HPD) intervals indicate that the *T_min_* and *T_opt_* for survival probability were higher and that the *T_max_* was lower under temephos treatments (Fig. 1B). In particular, the *T_min_* of juvenile survival probability increased by ∼1.1°C for every 0.001 ppm increase in temephos, leading to profound differences a *T_min_* 14.4°C higher than baseline when exposed to the highest tested temephos treatment. The *T_max_* decreased by ∼0.4°C for every 0.001 ppm increase in temephos, leading to ∼3.1°C-5.6°C lower *T_max_’s* under all temephos treatments. In line with this uneven reduction in thermal limits, the *T_opt_* of juvenile survival probability increased with temephos, with the *T_opt_* being 4.4°C higher when the mosquitoes were exposed to the highest temephos treatment. Overall survival (height of curve) decreased with increasing temephos concentrations. Peak survival (height of the curve at *T_opt_*) was 52.2% the control when exposed to 0.007 ppm temephos, 41.4% when exposed to 0.009 ppm temephos, 31.3% when exposed to 0.0111 ppm temephos, and 24.6% when exposed to 0.0127 ppm temephos (Fig. 1A).

**Figure 1.**
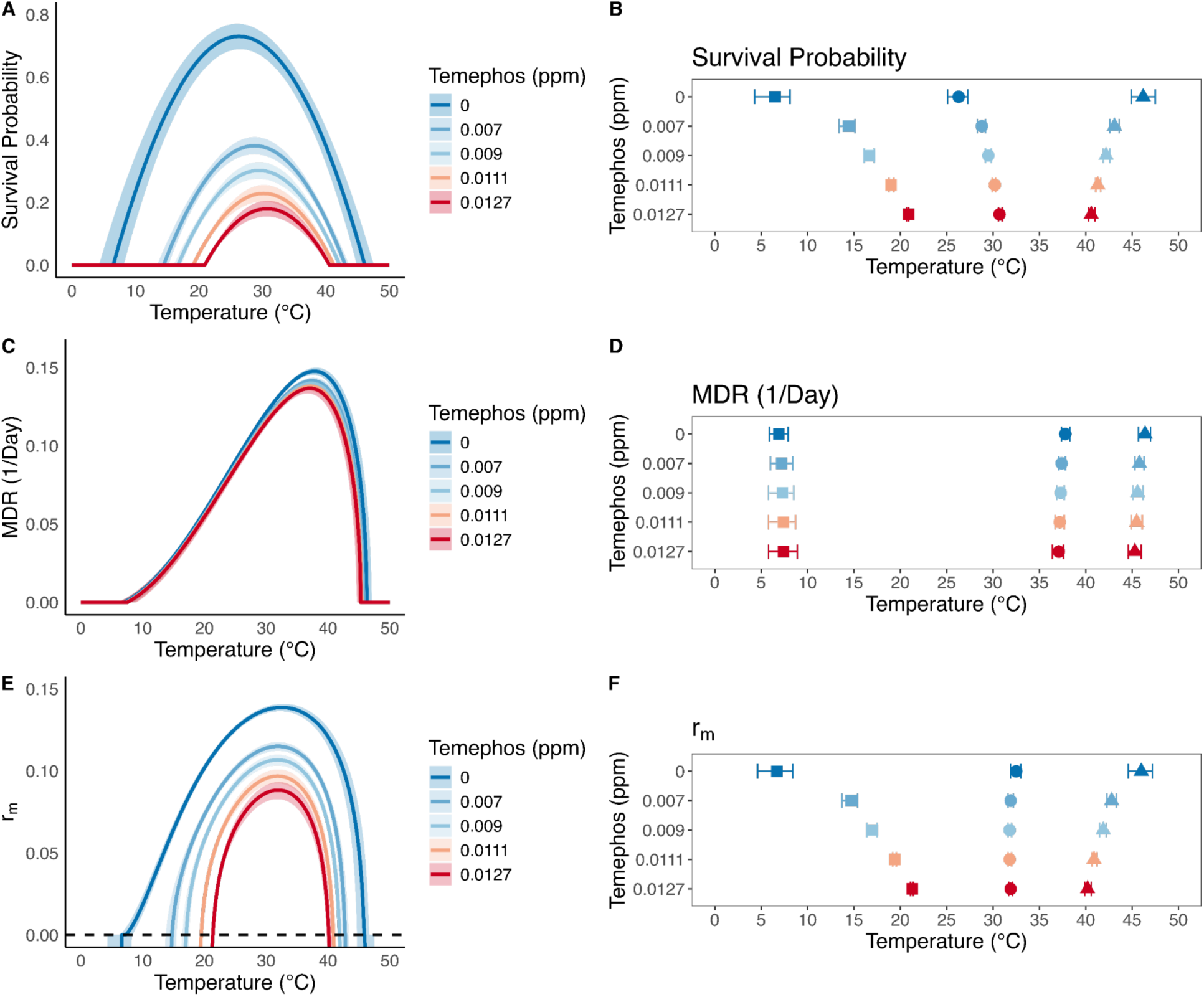
Effects of temephos and temperature on *Aedes aegypti* juvenile survival (A-B), development rate (MDR; C-D), and population growth rate (r_m_, E-F). Prediction bounds in **A**, **C**, and **E** are 95% credible intervals and prediction bounds in **B**, **D**, and **F** are highest posterior density (HPD) intervals each estimated from posterior distributions at each temephos level. In **B**, **D**, and **F**, the squares represent the thermal minimums (*T_min_*), the circles represent the thermal optimums (*T_opt_*), and the triangles represent the thermal maximums (*T_max_*).

### Effects of temperature and temephos concentration on juvenile development rate

Overlapping 95% HPD intervals indicated no differences among the *T_min_, T_max_,* and *T_opt_* of juvenile development rate among treatments and the control (Fig. 1C, 1D). In the control curve, the *T_opt_* of juvenile development rate was 37.8°C (37.4-38.3 95% HPD), the *T_min_* was 6.9°C (5.8-7.8 95% HPD), and the *T_max_* was 46.4°C (45.7-47.0 95% HPD). We observed a slight negative trend of −0.008 (−0.104 to −0.010 95% credible interval) with increasing temephos treatments. While this did not meaningfully influence the *T_max_*, the peak performance of juvenile development rate (height of the curve) was 4.1%-7.4% less than the control in our temephos treatments (Fig. 1C).

### Effects of temperature and temephos concentration on wing length

We observed a negative relationship between temperature and overall wing length of emerged mosquitoes (−0.028 ± 0.004 SE; p = 3.2e-14). No significant effect was observed for temephos concentration or the interaction between temephos concentration and temperature for overall wing length. We found similar results when analyzing just the female mosquitoes that emerged. We observed a negative relationship between temperature and female wing length (−0.03 ±0.005 SE; p = 1.28e-09), but no significant effect for temephos concentration or the interaction between temephos concentration and temperature.

### Effects of temperature and temephos concentration on population growth rate

Across temephos concentrations, *r_m_* exhibited a unimodal relationship with temperature (Fig. 1E). In our control curve, *r_m_* tended to be positive between 6.7°C (4.8°C-8.4°C) and 46.0°C (44.6°C-47.2°C). In our treatment groups, the minimum temperature for positive *r_m_* increased by ∼8.0°C-14.6°C, and the maximum temperature decreased by ∼3.2°C-5.8°C (Fig. 1F). Despite significant shifts in the thermal limits, the optimum temperature for positive *r_m_* was not different between the control and treatment groups. Peak *r_m_* (height of the curve at the optimum) was 82.9% the control when exposed to 0.007 ppm temephos, 76.9% when exposed to 0.009 ppm temephos, 69.8% when exposed to 0.0111 ppm temephos, and 63.6% when exposed to 0.0127 ppm temephos (Fig. 1E).

### Mapping Thermal Suitability

In the temperature alone predictions (Figs 2A, D, G, J), high *r_m_* values of *Ae. aegypti* were consistently predicted in the low elevation zones of the tropics, regardless of time of year, while low *r_m_* values were consistently predicted in cooler regions in space or season. Cold regions in Alaska, northern Canada, Greenland, and northern Russia never experienced conditions that favored positive population growth; neither did high-elevation regions such as the Himalayas or the Andes. Temporally, regions with high seasonal variation in temperature saw substantial swings in *r_m_*. For example, the midwestern United States was predicted to have positive *r_m_* from April to September but negative *r_m_* from October through March. We saw opposite temporal patterns in the Southern Hemisphere, with regions in southern Argentina and Australia having positive *r_m_* from October through March and negative *r_m_* from April through September.

**Figure 2.**
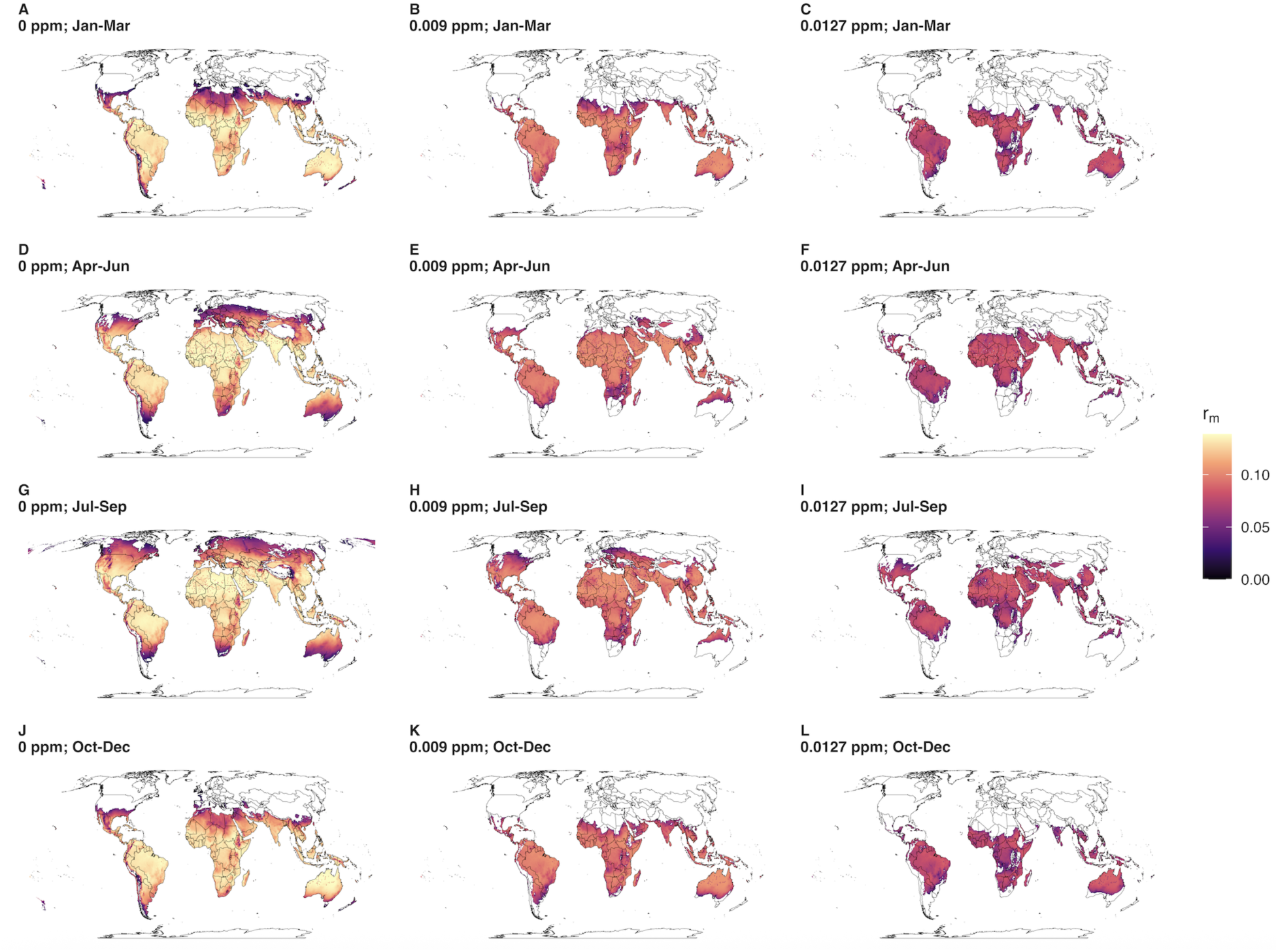
Spatial and temporal patterns of *Aedes aegypti* population growth rate (r_m_) based on mean seasonal temperature in 2024 (A-C: January through March; D-F: April through June; G-I: July through September; J-L: October through December) and temephos (A, D, G, I: 0 ppm; B, E, H, K: 0.009 ppm; C, F, I, L: 0.0127 ppm). White indicates areas with a negative r_m_. These maps serve as a heuristic tool alone and should not be interpreted as species distribution predictions.

For the temperature and temephos predictions (Figs 2B-C, E-F, H-I, K-J), temephos exposure decreased the *r_m_* of *Ae. aegypti* across all mapped locations. As we observed the largest effect of temephos on *r_m_* at low temperatures, it is not surprising that the largest differences in global predictions were observed in areas that were cooler in space and time. In the Northern Hemisphere, the greatest spatial restrictions of *r_m_* by temephos occurred in the United States, Mexico, Europe, and central Asia. In the Southern Hemisphere, the greatest predicted spatial restrictions of *r_m_* by temephos were seen from April-September, particularly in southern South America, southern Africa, and Australia. In Africa, pockets of shifts from positive to negative *r_m_* with temephos exposure were observed in high elevation regions, such as the Ethiopian Highlands and western Tanzania. As we lack high resolution on insecticide use, exposure, and resistance globally, these maps serve as heuristic illustrations of the potential effects of insecticide exposure on the thermal suitability of *Ae. aegypti*.

## Discussion

Our results demonstrate that increasing temephos exposure can not only reduce the population growth rate of *Ae. aegypti* mosquitoes but also reduce its thermal breadth. These patterns were driven by the interactive effect of temephos exposure and temperature on juvenile survival. While increasing temephos decreased juvenile survival across temperatures, it disproportionately reduced survival at cooler temperatures resulting in upward shifts in the *T_opt_* and *T_min_*. These results have three important implications. First, they provide further evidence that temperature does not act independently on mosquito traits. Second, they indicate that temephos is most effective at reducing mosquito population growth at lower temperatures despite lower expected toxicity. Third, they identify that insecticide control may become more challenging in warming environments.

The temperature-dependent patterns in juvenile survival and juvenile development rate were broadly consistent with previous studies demonstrating unimodal TPCs (Mordecai et al., 2019), and the linear decrease in wing length with temperature was consistent with general arthropod temperature-size theory (Atkinson, 1995; Klok & Harrison, 2013) and mosquito literature (Huxley et al., 2022; Mohammed & Chadee, 2011). Regardless of temephos concentration, we found non-linear, unimodal curves to adequately describe the patterns. However, under control conditions, we estimated broader thermal limits than those reported in prior meta-analyses (Mordecai et al., 2017, 2019), with lower *T_min_* and higher *T_max_* values for both survival and development rate. In contrast, *T_opt_* estimates were similar to previously published values. These differences likely reflect methodological and population-level variation, as Mordecai et al. (2017, 2019) synthesized data across multiple mosquito populations and experimental conditions, whereas our estimates are derived from a single field population. Population-level variation in mosquito thermal performance is well documented and underscores the importance of locally parameterized models.

The reduction in juvenile survival as temephos exposure increases is consistent with previous studies demonstrating concentration-dependent mortality in *Ae. aegypti* (Palomino et al., 2022; Tikar et al., 2009). In contrast, sublethal effects of temephos on the juvenile development rate are often weak or absent in previous studies (Mpho et al., 2001; Muturi et al., 2011; Wiwatanaratanabutr & Kittayapong, 2006), similar to our findings. Reported effects of temephos on the wing length, however, are more variable. While some studies have demonstrated no effect (Wiwatanaratanabutr & Kittayapong, 2006), others have shown that larval exposure results in slightly larger adults upon emergence (Mpho et al., 2001; Muturi et al., 2011). The magnitude of these effects is generally small, and given reduced sample sizes at high temperatures and concentrations, our study may have been underpowered to detect subtle differences. Finally, the decline in predicted population growth (*r_m_*) with increasing temephos exposure is not surprising given the strong reductions in juvenile survival, a primary determinant of fitness.

Beyond the main effects of temperature and temephos exposure, our results demonstrate that temephos exposure reshapes the thermal performance of juvenile survival and, consequently, the predicted population growth of *Ae. aegypti*. While temephos decreased juvenile survival across temperatures, these effects were strongest at cooler temperatures, resulting in an upward shift in thermal performance and a narrowing of thermal breadth. This response suggests that temperature modifies insecticide toxicity asymmetrically rather than uniformly increasing mortality across conditions.

We propose that temperature–temephos interactions arise from distinct mechanisms at low versus high temperatures. At low temperatures, slowed metabolism and development (Mordecai et al., 2019; Schulte, 2015) can increase exposure time to insecticides and decrease availability of detoxification enzymes (Lalouette et al., 2011), greatly reducing survival. In contrast, elevated metabolic activity and upregulation of detoxification enzymes with warming temperatures (Durak et al., 2021; Kumar et al., 2023; Neven, 2000) may partially buffer toxicity through enhanced detoxification efficiency (Agyekum et al., 2022). A tipping point is reached near the critical thermal maximum, however, as the energetic demands of coping with thermal stress may compromise detoxification capacity, leading to synergistic mortality under concurrent high temperature and insecticide exposure (González-Tokman et al., 2020). While no study has evaluated insecticide-temperature interactions across the full thermal breadth of mosquitoes, our findings are inconsistent with Muturi et al (2011), who found that a temperature-malathion interaction reduced survivorship in *Ae. albopictus* and *Culex restuans* most at 25°C and 30°C, but not at 20°C, suggesting that cool temperatures buffer against negative pesticide effects. Our results are also inconsistent with Mpho et al (2001), who found no significant temperature-temephos interaction on these traits in *Culex quinquefasciatus*. Our results could differ from these studies due to differences in mosquito species, insecticides, and/or temperature ranges, particularly the inclusion of cooler temperatures where we observed the largest effects.

Due to the effects of temperature and insecticide exposure on juvenile survival, temephos increased the thermal minimum, decreased the thermal maximum, and reduced overall thermal breadth for mosquito *r_m_*. This, in turn, led to distinct spatial and seasonal predictions for the predicted *r_m_* compared to predictions considering the effect of temperature in isolation. The largest insecticide-induced narrowing of the predicted thermal range of *r_m_* occurred in cooler regions. For current mosquito control, these results suggest that insecticides may be most effective in cooler regions and at cooler times of year. These results also demonstrate that many regions across the globe (particularly those that *Ae. aegypti* are invasive to) are currently below the thermal optima for *r_m_*. While insecticides may be particularly effective now, continued warming could shift these regions closer to the thermal optima, compromising the success of these control efforts. Together, these results highlight the importance of incorporating pesticide exposure into climate suitability and risk models for mosquito-borne disease.

Several limitations should be considered when interpreting these findings. First, low survival at high temperatures and temephos concentrations reduced statistical power to detect treatment differences in *T_opt_* and *T_max_* for juvenile development rate. Although we observed trends towards reduced development at the highest temephos concentration (0.0127 ppm), larger sample sizes would be required to determine whether these differences are robust.

Second, our experimental design did not account for field-relevant heterogeneity such as diurnal temperature fluctuations, which may alter mosquito physiology and temephos decay and toxicity. While constant temperatures simplify the natural environment, they are a standard first step in isolating the impact of temperature on performance (Sinclair et al., 2016), and mean temperatures have recently been demonstrated to better predict thermal limits of mosquito-borne disease transmission than rate summation of diurnal patterns (Shocket et al., 2025). Extending these results to diurnal temperature fluctuations remains an important area of future work, however.

Third, we did not account for population-level variation in insecticide resistance (Moyes et al., 2017). Resistance may have important implications for two reasons: (i) resistant mosquitoes are more likely to be exposed to a sublethal concentration of an insecticide, and (ii) resistance tends to come with fitness costs, which may influence the shape of the temperature- and insecticide-dependent performance curves. Characterizing how resistance influences the temperature-by-insecticide interaction on *Ae. aegypti* aquatic stages represents an important avenue for future research. Finally, our calculation of population growth rate assumes no carry over effects of temephos on adult mosquito fitness and does not account for variation in environmental carrying capacity for *Ae. aegypti* (Sanil & Shetty, 2012). For this study, our goal was to isolate the effects of temperature and temephos on the juvenile traits of a metric of population fitness. Thus, interpretation of the mapped population growth rate should not extend to realized *Ae. aegypti* distribution and abundances. Despite these limitations, this study demonstrates that chemical control can modify TPCs, reinforcing evidence from other systems (e.g., Nguyen et al [ 2021]) that also show that chemical control has the strongest effects at cooler temperatures, potentially suggesting a general pattern.

In conclusion, our results demonstrate that insecticide exposure can reshape TPCs rather than simply reduce population growth uniformly across temperatures. Even low concentrations of temephos altered the thermal limits and optimum of *Ae. aegypti* juvenile survival and population growth, with implications for predicting control efficacy across space and time. More broadly, this study highlights the need to integrate anthropogenic stressors with climatic drivers when optimizing pest management strategies under changing environments. Future research should evaluate whether similar temperature-dependent effects occur for other insecticides, resistant mosquito populations, and traits directly linked to arbovirus transmission risk.

