## Supplementary Information for "Insecticide temephos alters thermal dependence of dengue vector"

**Contents:**

**Table S1.** Response surface treatment design.

**A. Mathematical models for Thermal Performance Curves**

**B. Sensitivity analyses results.**

**Figure S1.** Sensitivity of intrinsic growth rate

**References for Supplementary Materials**

**Table S1. Treatments tested to estimate survival probability, development rate, and wing length in juvenile *Aedes aegypti* mosquitoes.** 'X' denotes a temperature-by-temephos treatment tested, and a blank cell denotes a treatment *not* tested.

| | | **Temperature (°C)** | | | | | | |
|---|---|---|---|---|---|---|---|---|
| **Temephos (ppm)** | | **17** | **21** | **25** | **29** | **33** | **37** | **41** |
| | **0** | **X** | **X** | **X** | **X** | **X** | **X** | **X** |
| | **0.007** | **X** | | | **X** | | | **X** |
| | **0.009** | **X** | **X** | **X** | **X** | **X** | **X** | **X** |
| | **0.0111** | **X** | | | **X** | | | **X** |
| | **0.0127** | **X** | **X** | **X** | **X** | **X** | **X** | **X** |

### A. Mathematical models for Thermal Performance Curves

*Quadratic model for juvenile survival probability*

For quadratic models used for fitting juvenile survival probability ($\rho EA$) trait data across temperatures ($T$), we implemented the following model in which $T_{min}$ and $T_{max}$ were monotonically dependent on temephos concentration ($C$). Priors for $b_a$ and $b_b$ (the $T_{max}$ and $T_{min}$ at 0 ppm temephos) were drawn from normal distributions centered on mean values extracted from Mordecai et al (2019) with σ's of 2.5. The priors for $m_a$ and $m_b$ (the slopes of how $T_{max}$ and $T_{min}$ change with temephos concentration) were drawn from a normal distribution centered at 0 with a σ of 1, allowing the slope to be positive or negative. The prior for $q$ was drawn from a normal distribution centered on mean value for $q$ and a σ equal to twice the standard error from Mordecai et al (2019). As we were modeling probabilities, the values were restricted to be between 0 and 1. The full mathematical model is as follows:

$$p_i = (-q)(T_i - T_{max_i})(T_i - T_{min_i})(T_{min_i} \leq T_i \leq T_{max_i})\, T(0, 1)$$

$$T_{max_i} = b_a + (m_a)(C_i)$$

$$T_{min_i} = b_b + (m_b)(C_i)$$

$$\rho EA_i = bin(p_i, n_i)$$

$$q \sim norm(0.00599, 0.000862)$$

$$b_a \sim norm(38.3, 2.5)$$

$$b_b \sim norm(13.6, 2.5)$$

$$m_a \sim norm(0, 1)$$

$$m_b \sim norm(0, 1)$$

*Brière model for juvenile development rate*

For Brière models used for fitting juvenile development rate (*MDR*) trait data across temperatures (*T*), we implemented the following model in which $T_{min}$ and $T_{max}$ were monotonically dependent on temephos concentration (*C*). Model formulation and priors were chosen using the same process as above. As the juvenile development rate cannot be negative, we restricted the values to only be positive. The full mathematical model is as follows:

$$\mu_i = (q)(T_i)(T_i - T_{min_i})\sqrt{(T_{max_i} - T_i)}(T_{min_i} \leq T_i \leq T_{max_i})$$

$$T_{max_i} = b_a + (m_a)(C_i)$$

$$T_{min_i} = b_b + (m_b)(C_i)$$

$$MDR_i = norm(\mu_i, \sigma)\, T(0, \infty)$$

$$q \sim norm(0.0000783, 0.0000214)$$

$$b_a \sim norm(39.1, 2.5)$$

$$b_b \sim norm(11.6, 2.5)$$

$$m_a \sim norm(0, 1)$$

$$m_b \sim norm(0, 1)$$

$$\sigma \sim uniform(0, 1)$$

### B. Sensitivity analyses results

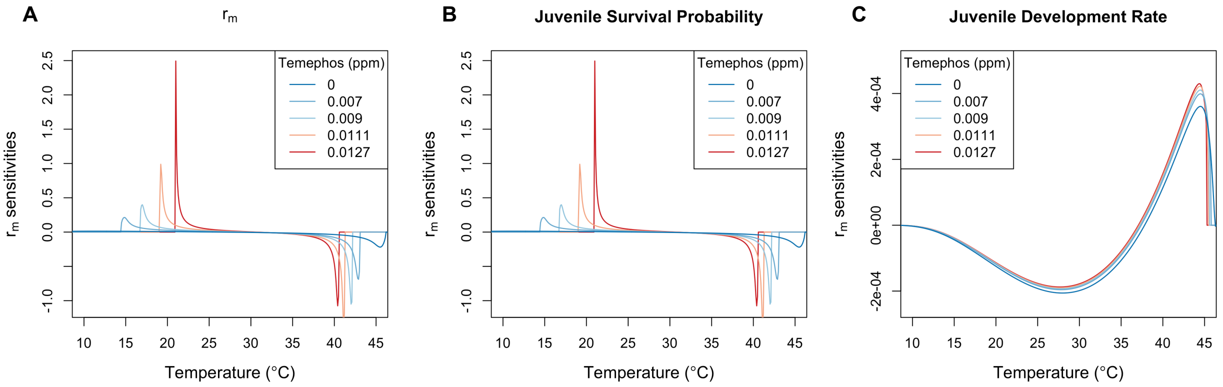


**Figure S1. Sensitivity of intrinsic growth rate, $r_m$, across temperatures and temephos concentrations to traits in juvenile *Aedes aegypti* mosquitoes. A** is the derivative of $r_m$ with respect to temperature, whereas **B** and **C** are the partial derivatives of juvenile survival probability and juvenile development rate with respect to temperature. Please note that **A** and **B** are on the scale from -1 to 2.5, whereas **C** is on the scale from -0.00025 to 0.0005.

Variation in $r_m$ across temperatures and temephos concentrations was almost entirely driven by juvenile survival probability (Fig. S1). Temephos exposure increased the sensitivity of $r_m$ and juvenile survival probability below 25°C and above 35°C. Sensitivity of juvenile development rate was minimal with the steepest sensitivities being above 35°C (the peak and decline of the Brière curve). These results emphasize that the $r_m$ is highly sensitive to the interaction between temperature and temephos, particularly due to strong effects on juvenile survival probability at low and high temperatures.
